# Transposon mutagenesis uncovers the genetic landscape of streptomycin susceptibility and implicates SbmA in aminoglycoside uptake in *Escherichia coli*

**DOI:** 10.64898/2026.08.18.745456

**Authors:** Weine J Kok, James Griffith, Damien Merke, Adam F Cunningham, Ian R Henderson, Emily C A Goodall

## Abstract

Aminoglycosides are critical antibiotics with partially elucidated mechanisms of uptake and action in Gram-negative bacteria. Importantly, although energy-dependent uptake across the inner membrane has been well-established, the specific molecular mechanisms involved have not been definitively identified. To deepen understanding of genetic factors influencing susceptibility and resistance to streptomycin, we applied transposon insertion sequencing in *Escherichia coli* K-12. This approach identified both known and novel genes whose disruption increased susceptibility, including those involved in respiration, protein export, cell division, and uncharacterised functions. Notably, voltage-sensitive membrane dye-based assays revealed that many susceptible mutants did not display inner membrane hyperpolarisation as often assumed. Conversely, disruption of certain genes, such as the inner membrane antimicrobial peptide transporter *sbmA*, conferred low-level resistance, with *sbmA* overexpression increasing streptomycin sensitivity, suggesting its role in aminoglycoside uptake. These findings refine the model of aminoglycoside interaction with various pathways and highlight potential targets for adjuvant therapies to combat antimicrobial resistance.

## Introduction

Bacterial antimicrobial resistance (AMR) has increased over the past decade. Between 1991 and 2021, annual AMR-attributable deaths in individuals aged 5 and over has increased by 35% (1). The most critical or threatening bacterial pathogens identified by the World Health Organisation (WHO) are diderms, such as Gram-negative bacteria or *Mycobacterium tuberculosis* (2). If left unaddressed, AMR-associated deaths are estimated to increase to 9-10 million by 2050 (1). With a drying drug pipeline, new strategies and treatment regimens with existing antibiotics classes are urgently needed.

Aminoglycosides are one of the major antibiotic classes currently in use in both clinical and veterinarian contexts. The uptake and mechanism of action of aminoglycosides have been extensively studied, particularly for Gram-negative species and the model *E. coli* (3, 4). Aminoglycosides are thought to first attach to the lipopolysaccharide (LPS) layer through polar interactions (5), then enter the cell through the outer membrane (OM) via OM porins OmpC and OmpF (5, 6). Next, the drug molecules traverse the inner membrane (IM) in a proton motive force (PMF)-dependent fashion (7). Whether or not proteins are involved in this step remains unclear, but recent molecular dynamic and biophysical studies of aminoglycoside interactions with bacterial biomimetic membranes suggest that aminoglycosides are unlikely to spontaneously traverse across the membrane, consequently suggesting uptake via protein channels is possible (8). Following entry into the cytoplasm, aminoglycosides bind to the A-site of 16S ribosomal subunits, causing tRNA mismatching and ultimately leading to the mistranslation of proteins. These mistranslated proteins are thought to insert into the IM, creating non-specific channels that facilitate further aminoglycoside entry and continued inhibition of the 16S ribosome (9). Finally, it has been proposed that aminoglycoside-induced reactive oxygen species, metabolic perturbations and/or membrane hyperpolarisation ultimately lead to cell death, although no conclusive evidence has been presented for each of these hypotheses (10–15).

To address gaps in the understanding of aminoglycoside uptake and mechanism of action, a number of functional genomics-based studies have been applied (16, 17). A collection of single gene knockout mutants in *E. coli* K-12, the Keio collection, were screened in sub-inhibitory concentrations of the aminoglycosides streptomycin, gentamicin and tobramycin (16, 17), where mutant fitness was assessed by colony size. This work identified known hypersusceptible mutants, including those from ATP synthase genes (*atpBEFHAGDC*) (18). However, such an approach can suffer from poor reproducibility (17), and be biased by secondary mutations (19). In another study, microarray expression profiles of *E. coli* K-12 cells treated with inhibitory concentrations of the aminoglycosides kanamycin and gentamicin identified changes in expression of protein secretion, envelope stress responses and central carbon metabolism pathways, which were validated with their respective knockout mutants (11). For example, aminoglycoside hypersensitivity was displayed in *secG* mutants, a component of the Sec translocon required for protein translation across the IM, as well for *hflC* and *hflK* mutants, which negatively regulate the protease FtsH responsible for degrading membrane-associated proteins (11). A more sensitive contemporary technique for such genetic screens is transposon insertion sequencing (TIS). TIS couples transposon mutagenesis with high-throughput sequencing to quantify gene fitness of virtually every non-essential gene in the query bacterial organism in a given condition (20–22). Furthermore, TIS overcomes the strict loss-of-function limitations of the Keio collection by assaying intergenic/non-annotated regions, thereby enabling the identification of gain-of-function phenotypes, such as the overexpression of downstream genes. In the context of antibiotic screens, TIS has been widely used to identify genes responsible for resistance and hypersensitivity to a particular antibiotic by subjecting the transposon mutant library to a sub-inhibitory or inhibitory concentration of antibiotic, elucidating genes that are essential for survival in that antibiotic (23–27). Recently, TIS has been applied to *E. coli* K-12 strains resistant to streptomycin, gentamicin, or neomycin (28). However, we reason that using strains with established resistance to aminoglycosides may obscure significant genetic determinants of susceptibility that would otherwise be identifiable in an aminoglycoside-sensitive genetic background.

Remarkably, 80 years after the first use of streptomycin, the first antibiotic in this class, major gaps in understanding the mechanism of action of aminoglycosides remain, including how aminoglycosides are trafficked through the IM and how they are tolerated by cells. Here, we utilised TIS to globally assess the genetic determinants for streptomycin susceptibility and resistance in *E. coli* K-12 as a model for other Gram-negative bacteria. We uncovered diverse pathways associated with ATP synthesis, two-component signal transduction systems, cell division, and IM transporters that were vulnerable to streptomycin stress when disrupted. Although many of these mutants utilise a proton gradient for function, their respective deletion mutants generally did not display increased IM polarisation, suggesting the presence of proton motive force (PMF)-independent pathways to tolerate aminoglycoside lethality. Additionally, we found numerous genes whose disruption conferred increased resistance to streptomycin, such as those belonging to respiratory complexes and their regulatory systems. Lastly, our data showed that overexpression of the IM proton symporter SbmA, which is widely conserved across diderm bacteria, resulted in hypersensitivity to streptomycin, and other aminoglycosides, suggesting a role for SbmA in facilitating PMF-dependent uptake of aminoglycosides. Taken together, our work provides a more nuanced model of how streptomycin, and more broadly aminoglycosides, exert their antimicrobial action.

## Results

### Identification of mutants susceptible to a sub-inhibitory dose of streptomycin

To gain a better understanding of how streptomycin exerts its antimicrobial activity, we exposed an *E. coli* K-12 transposon library to sub-inhibitory concentrations of streptomycin. At a sub-inhibitory concentration, we expect that mutants sensitive to streptomycin will be outcompeted during growth and therefore less abundant within the overall pool, as we have demonstrated previously for polymyxin B (27), while neutral gene-disruption events will have a negligible impact. Conversely, we hypothesised that mutants that are more fit will proliferate and increase in relative abundance in the overall mutant pool, including those encoding for IM/OM putative aminoglycoside transporters or complexes generating the energetics (i.e. PMF) necessary for aminoglycoside uptake. We identified a suitable concentration of streptomycin for selection through growth assays of the parent *E. coli* K-12 strain BW25113 in varying concentrations of streptomycin (**S. Fig. 1A**). We identified 16 µg/ml as the minimal inhibitory concentration (MIC), defined here as the lowest concentration required to inhibit growth in LB media. For our transposon library screen, we selected a concentration of 4 µg/ml (1/4x MIC), as this was the highest concentration that did not significantly inhibit the growth rate of *E. coli* BW25113, allowing for the identification of mutants hypersensitive to streptomycin.

**Figure 1.**
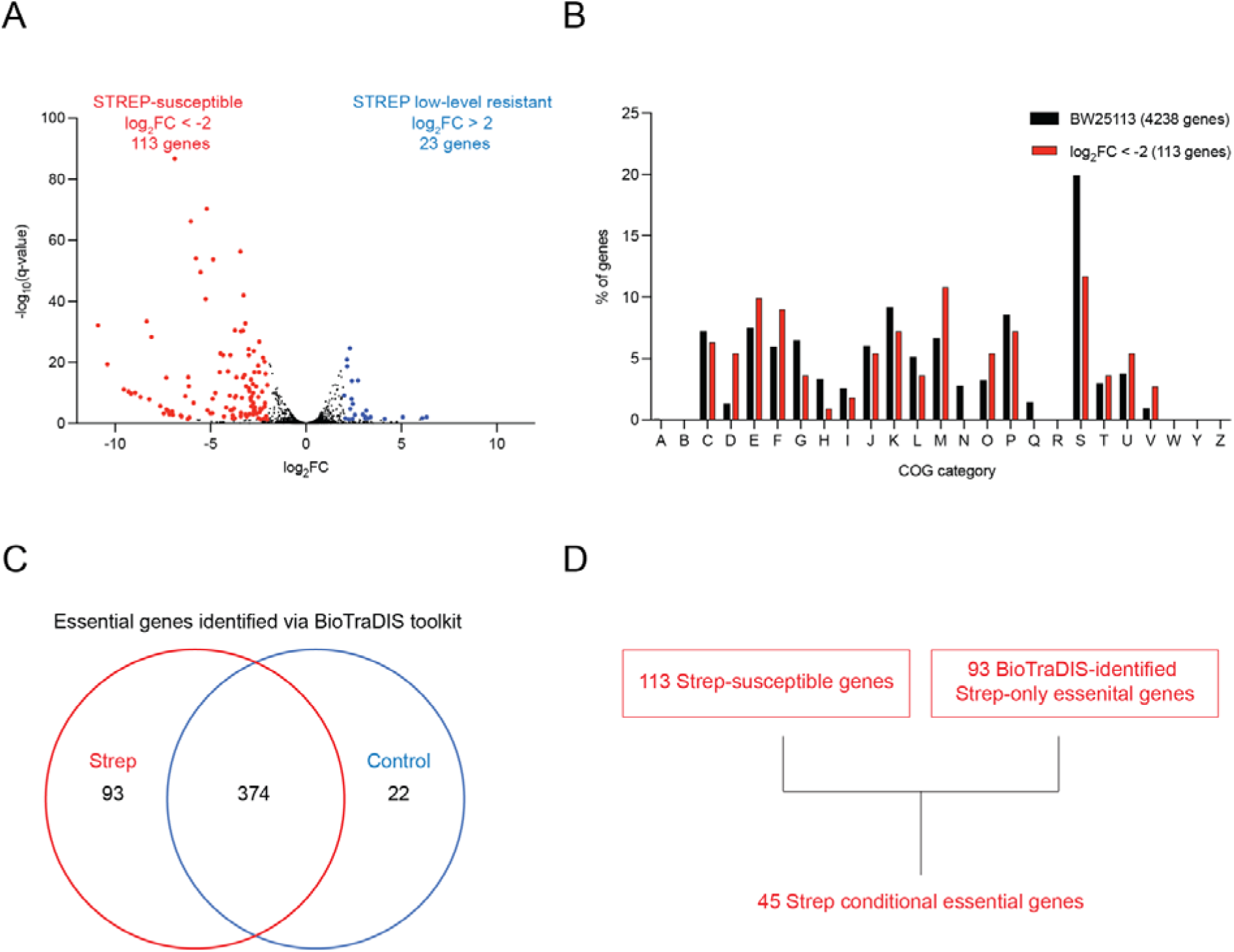
Identification of conditionally essential genes required for surviving streptomycin stress. **(A)** Volcano plot depicting gene fitness of *E. coli* BW25113 transposon mutant library grown in a sub-inhibitory concentration of streptomycin (Strep; 4 μg/mL). Mutants that have a lower gene fitness score (log_2_FC ≤ −2) than the untreated control are deemed Strep-susceptible (in red), whereas mutants that are fitter than the control (log_2_FC ≥ 2) are deemed STREP-resistant (in blue). All significant gene hits all have an q-value ≤ 0.05. **(B)** Cluster of Orthologous Groups (COG) analysis of gene hits. **(C)** Venn diagram depicting unique or shared essential genes identified from Step-treated or control libraries using the tradis_essentiality package from the BioTraDIS toolkit. **(D)** Genes were identified as conditionally essential in streptomycin by meeting two criteria. A conditionally gene was required to **(i)** have at least a four-fold lower transposon insertion count in the streptomycin-treated library compared to the control, and **(ii)** be classified as uniquely essential by from the BioTraDIS toolkit in the streptomycin-treated library.

The success of TIS experiments is highly dependent on the mutant diversity or density of the input transposon mutant library (29). Previously, we reported a dense transposon mutant library in the *E. coli* K-12 strain BW25113, comprising nearly 1 million unique mutants, each with a randomly inserted mini-Tn*5* transposon carrying a chloramphenicol resistance marker (30). We chose this library for our antibiotic selection screen as it is a highly saturated library that has been well characterised. We grew the transposon library in LB supplemented with 4 μg/mL (1/4x MIC), across three passages of culture, each growing from an OD_600_ of 0.05 to 0.5, in duplicate (**S. Fig. 1B**), and included an LB-only control for comparison. Three passages within this OD_600_ range allowed cells to grow through ∼9-10 generations, as well as maintaining them within the exponential phase of growth critical for aminoglycoside activity (31). Cells were harvested and the transposon insertion sites identified through sequencing of the transposon-genomic DNA junctions using the Illumina sequencing platform. Overall, we recovered approximately 2 M reads per replicate that were mapped to the *E. coli* BW25113 reference genome (EMBL accession: CP009273; S. Table 1). Replicates were highly correlated (**S. Fig 1C**), and these data were therefore pooled (per condition) to visualise total transposon insertion sites *en masse*. On a genome-wide view, the insertion profiles arising from each condition were sinusoidal, with insertion read depth peaking around the 3.7 Mb mark corresponding with the chromosomal replication origin (**S. Fig. 2**). This origin-proximal bias is likely attributable to multifork replication (29, 32, 33). Since this bias was noted in both the control and streptomycin-treated libraries, we did not correct for it.

**Figure 2.**
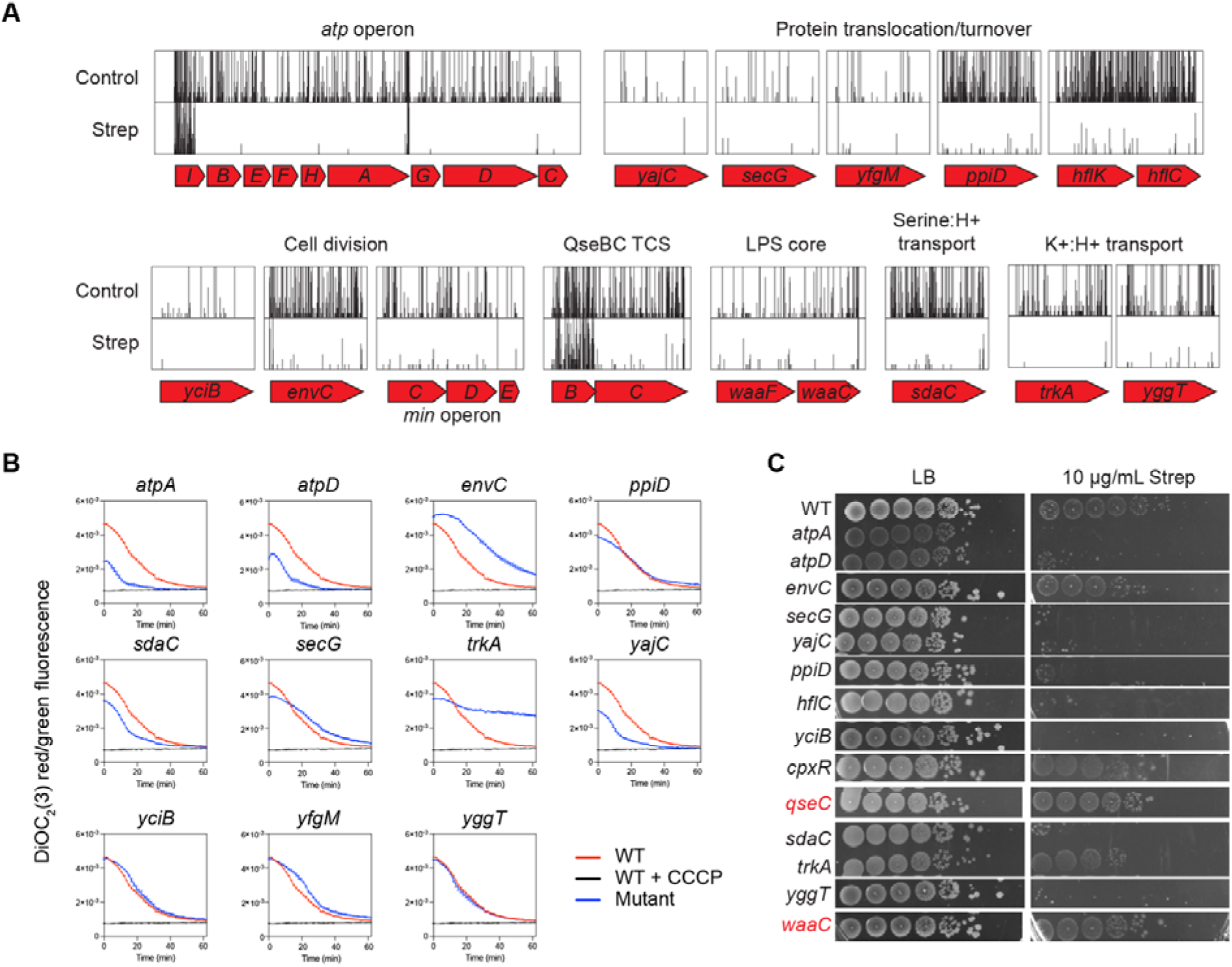
Validation of streptomycin-susceptible mutants. **(A)** Transposon insertion plots for the control and Strep outgrowth conditions of selected genes identified to be susceptible to Strep when disrupted. The vertical axes of the insertion plots represent normalised insertions (in counts per million), with both control and Strep plots axis heights set to a maximum of 10. **(B)** Membrane potential assay of *E. coli* BW25113 wildtype or respective deletion mutants. Cells were grown to exponential phase (OD_600_ = 0.4) in LB, normalised, washed with PBS + EDTA+ glucose, and stained with DiOC_2_(3). Redshifted (Ex: 488 nm Em: 650 nm) and green (Ex: 488 nm Em: 560 nm) fluorescence were measured in a 96-well plate reader over 60 min. Reported values shown are expressed as a ratio of red/green emission values. As a depolarised control, WT cells were incubated with the protonophore CCCP which is expected to dissipate the PMF. Experiments were performed with three biological replicates, and error bars indicate SEM. **(C)** Overnight cultures were normalised to OD_600_ = 0.1, ten-fold serially diluted, before being inoculated onto LB agar supplemented with 10 µg/ml streptomycin. Mutants that were not sensitive to streptomycin are highlighted in red.

To identify mutants that were susceptible (less fit) or more fit (henceforth termed low-level resistant/tolerant) in sub-inhibitory concentrations of streptomycin, we used the BioTraDIS tradis_comparison.R script (34), which implements the edgeR package to measure changes in read depth (per gene) between conditions. The number of reads mapped to each gene is used as a proxy for mutant abundance. Overall, we identified 113 streptomycin-susceptible genes (defined as log_2_FC ≤ −2, q-value ≤ 0.05) and 23 low-level resistant/tolerant genes (defined as log_2_FC ≥ 2, q-value ≤ 0.05) (**Figure 1A**). We reviewed the Cluster of Orthologous Groups (COG) functional classification of the 113 streptomycin-susceptible genes and found that they were largely enriched in the ‘S’ (function unknown), ‘M’ (cell envelope biogenesis), ‘E’ (amino acid transport and metabolism) and ‘F’ (nucleotide transport and metabolism) categories (**Figure 1B**).

Our 113 streptomycin-susceptible genes identified via edgeR (tradis_comparison.R) could contain false positives in genes with few unique insertions, as sufficient read counts at those scarce sites can still produce statistically significant fold-change values, warranting a complementary analysis. Therefore, we re-analysed the TIS data using the tradis_essentiality tool from BioTraDIS to define essential genes from the LB-only and streptomycin-treated TIS libraries (see Methods). Intersecting both essential gene lists yielded 93 essential genes unique to the treated library (**Figure 1C**). Manual inspection of the transposon insertion profiles of these 93 streptomycin-unique essential genes found many of these genes contained sparse insertions in the control. To filter these genes, we compared the data from the edgeR streptomycin-susceptible genes and BioTraDIS tradis_essentiality streptomycin-unique essential genes and selected to only study genes common to both outputs, as we hypothesised that those genes should produce the strongest streptomycin sensitivity phenotype. This approach revealed 45 genes common to both gene lists, which we defined here to be streptomycin conditionally essential (**Figure 1D**) (Table 1). We further refined our data by manual inspection of transposon insertion profiles in these genes and found that only *ftsL* and *minE* had sparse insertions in the control condition. Indeed, *ftsL* and *minE* were deemed essential in the Keio collection (35). Given the relatively small proportion of hits that were falsely identified, our analysis approach highlights its accuracy in identifying hits associated with increased streptomycin sensitivity.

**Table 1.** Table of 45 streptomycin conditionally essential genes (i.e. possess log_2_FC ≤ −2, q-value ≤ 0.05 and deemed streptomycin-essential by BioTraDIS)

| Gene | Function | $\log_2FC$ | logCPM | q-value |
| --- | --- | --- | --- | --- |
| *atpD* | F1 sector of membrane-bound ATP synthase, beta subunit | -10.90 | 6.19 | 7.82E-33 |
| *atpB* | F0 sector of membrane-bound ATP synthase, subunit a | -10.41 | 5.74 | 4.22E-20 |
| *atpE* | F0 sector of membrane-bound ATP synthase, subunit c | -9.55 | 4.99 | 7.26E-12 |
| *yciB* | IspA family inner membrane protein | -9.30 | 4.73 | 3.77E-11 |
| *atpH* | F1 sector of membrane-bound ATP synthase, delta subunit | -9.18 | 4.63 | 1.53E-10 |
| *pgm* | phosphoglucomutase | -8.95 | 4.37 | 8.13E-11 |
| *pstB* | phosphate transporter subunit | -8.67 | 4.13 | 1.96E-09 |
| *atpA* | F1 sector of membrane-bound ATP synthase, alpha subunit | -8.35 | 6.26 | 3.65E-34 |
| *atpF* | F0 sector of membrane-bound ATP synthase, subunit b | -8.22 | 3.71 | 1.12E-08 |
| *trkA* | NAD-binding component of Trk potassium transporter | -8.09 | 6.01 | 4.90E-29 |
| *sapF* | antimicrobial peptide transport ABC system ATP-binding protein | -7.63 | 3.19 | 1.70E-06 |
| *atpG* | F1 sector of membrane-bound ATP synthase, gamma subunit | -7.32 | 5.32 | 1.01E-15 |
| *hns* | global DNA-binding transcriptional dual regulator H-NS | -7.31 | 2.98 | 2.27E-05 |
| *gmhB* | D,D-heptose 1,7-bisphosphate phosphatase | -7.25 | 3.00 | 8.39E-05 |
| *sapC* | antimicrobial peptide transport ABC transporter permease | -7.24 | 2.79 | 6.13E-05 |
| *minE\** | cell division topological specificity factor | -7.01 | 2.71 | 1.29E-04 |
| *hflC* | modulator for HflB protease specific for phage lambda cII repressor | -6.88 | 7.86 | 2.04E-87 |
| *atpC* | F1 sector of membrane-bound ATP synthase, epsilon subunit | -6.32 | 4.33 | 1.18E-09 |
| *icd* | e14 prophage; isocitrate dehydrogenase, specific for NADP+ | -6.19 | 1.74 | 3.93E-02 |
| *ptsN* | sugar-specific enzyme IIA component of PTS | -6.16 | 5.58 | 7.10E-16 |
| *secG* | preprotein translocase membrane subunit | -6.14 | 5.04 | 6.10E-13 |
| *epmA* | Elongation Factor P Lys34 lysyltransferase | -5.89 | 3.93 | 1.52E-07 |
| *ppiD* | periplasmic folding chaperone, has an inactive PPIase domain | -5.52 | 7.37 | 2.75E-50 |
| *envC* | activator of AmiB,C murein hydrolases, septal ring factor | -5.25 | 7.33 | 1.64E-41 |
| *marB* | mar operon regulator, periplasmic | -5.01 | 3.37 | 4.72E-04 |
| *ftsL\** | membrane bound cell division protein at septum containing leucine zipper motif | -4.91 | 3.26 | 4.23E-04 |
| *sapB* | antimicrobial peptide transport ABC transporter permease | -4.89 | 4.41 | 7.80E-09 |
| *waaC* | ADP-heptose:LPS heptosyl transferase I | -4.75 | 5.18 | 9.60E-11 |
| *qseC* | quorum sensing sensory histidine kinase in two-component regulatory system with QseB | -4.34 | 6.69 | 4.28E-23 |
| *hldD* | ADP-L-glycero-D-mannoheptose-6-epimerase, NAD(P)-binding | -4.18 | 4.90 | 5.51E-06 |
| *pstS* | periplasmic phosphate binding protein, high-affinity | -4.09 | 6.18 | 7.89E-18 |
| *epmB* | EF-P-Lys34 lysylation protein; weak lysine 2,3-aminomutase | -3.95 | 4.97 | 4.68E-05 |
| *smpB* | trans-translation protein | -3.83 | 3.02 | 1.78E-03 |
| *yajC* | SecYEG protein translocase auxillary subunit | -3.77 | 4.49 | 1.22E-05 |
| *tolC* | transport channel | -3.73 | 7.08 | 3.24E-31 |
| *gor* | glutathione oxidoreductase | -3.67 | 5.87 | 1.00E-09 |
| *purA* | adenylosuccinate synthetase | -3.65 | 6.60 | 7.98E-16 |
| *rbfA* | 30s ribosome binding factor | -3.41 | 3.23 | 4.48E-03 |
| *acnB* | bifunctional aconitate hydratase 2/2-methylisocitrate<br>dehydratase | -3.41 | 4.49 | 3.14E-04 |
| *pstC* | phosphate transporter subunit | -3.36 | 5.70 | 3.46E-09 |
| *sapD* | antimicrobial peptide transport ABC system ATP-binding protein | -3.17 | 3.50 | 1.08E-03 |
| *rpoN* | RNA polymerase, sigma 54 (sigma N) factor | -3.11 | 4.90 | 6.11E-03 |
| *ftsX* | inner membrane putative ABC superfamily transporter permease | -2.96 | 3.93 | 8.39E-04 |
| *polA* | fused DNA polymerase I 5'->3' polymerase/3'->5'<br>exonuclease/5'->3' exonuclease | -2.91 | 6.14 | 3.60E-09 |
| *glnD* | uridylyltransferase | -2.03 | 4.97 | 4.94E-02 |

\*Gene was deemed essential in the Keio collection.

Among 45 streptomycin conditionally essential genes, we found mutants in the *atp* operon, which encodes for subunits of the ATP synthase, to be sensitive towards streptomycin (**Figure 2A**). Loss-of-function mutations in the *atp* operon have previously been reported to confer hypersensitivity to aminoglycosides, thus confirming the validity of our screen (4, 18, 36). Previously, *atp* mutants (*unc*) possessing compromised ATP synthase activity displayed increased streptomycin sensitivity and uptake (18).

Genes belonging to the Sec protein translocon, *yajC* and *secG,* and accessory Sec translocon genes, *yfgM*, *ppiD*, *hflC* and *hflK* were all found to be conditionally essential, consistent with previous reports (**Figure 2A**) (11, 28). PpiD acts as a periplasmic chaperone that assists in the maturation of OMP proteins (37, 38). HflC and HflK form a membrane complex with the protease FtsH (an essential protein), acting as a quality control complex and stress response regulator for cytoplasmic and IM proteins via proteolysis in *E. coli* (39), including stalled Sec complexes (40), and presumably aminoglycoside-induced mistranslated proteins.

Several of our hits were related to cell division, including *envC*, a peptidoglycan amidase regulator (41), and *minCDE*, a cell division regulatory complex that directs cell septa to exclusively form at the mid-cell (42, 43) (**Figure 2A**). Additionally, a gene of unknown function, *yciB*, was identified in our screen. YciB was previously reported to physically interact with several cell division and cell shape proteins, including RodZ, MurG and MreD (44).

Other hits included genes associated with LPS core oligosaccharide biosynthesis. From the streptomycin-treated library, we observed a depletion of insertions in the *waaC* and *waaF* genes. Both genes encode for heptosyltransferases that attach the first and second heptose sugars in the Kdo_2_ moiety of the inner LPS core respectively (45). Disruption of either gene would give rise to cells with truncated LPS (46). Mutants lacking either of these genes were previously shown to possess increased OM permeability to a variety of antibiotics, including aminoglycosides (47–49).

Members of several two-component systems were also implicated in streptomycin sensitivity when disrupted. Mutants of the transcriptional regulator of the CpxAR system, *cpxR,* were depleted relative to the total pool following exposure to streptomycin. The Cpx system is activated during IM stress, where a key response is to regulate the expression of respiratory complexes, which could in turn confer protection against aminoglycosides (50). Accordingly, loss-of-function mutations in *cpxR* were previously shown to confer a hypersensitive phenotype against aminoglycosides (11). In addition to the Cpx system, the sensor histidine kinase of the QseBC two-component system (TSC), *qseC,* was also identified. QseBC has been implicated in the regulation of flagellar motility, biofilm-formation, virulence factors and host immune resistance genes (51).

Lastly, we found several genes encoding IM import-associated proteins, such as *sdaC*, a serine:H+ importer (52, 53), and *trkA*, a NAD+ binding component of the TrkG/H potassium:H+ symporters (54, 55), that were indispensable under streptomycin stress. Interestingly, a *trkA* null mutation was reported to depolarise the IM potential, in part due to the de-regulation and subsequent opening of *trkH* channels to spontaneously uptake sodium ions into the cell (55). Related to TrkG/H, we also identified a putative potassium transporter gene, *yggT,* whose disruption conferred a fitness defect in streptomycin. This gene was found to be a multicopy suppressor for fitness defects in *E. coli* cells lacking the major potassium uptake systems (including TrkG/H) (56).

A caveat in our approach to identify streptomycin conditionally essential genes is that the BioTraDIS essentiality analysis sets an arbitrary insertion index score (IIS) threshold for identifying essential genes. This threshold is derived from the bimodal distribution of the IISs but inevitably forces the binary classification (essential vs non-essential/ambiguous) of genes. Such an approach could miss streptomycin-susceptible (i.e. log_2_FC ≤ −2) genes whose IIS were shy of those thresholds to be deemed as ‘essential’. For example, in our streptomycin-treated library, this threshold was an IIS of 0.0115; genes with an IIS < 0.0115 were deemed as ‘essential’. However, streptomycin-susceptible genes such as *sdaC* (serine:proton symporter; log_2_FC = −5.8, q-value = 9.16 x 10^−55^) and *yggT* (log_2_FC = −4.5, q-value = 1.16 x 10^−23^) had IISs of 0.0116 and 0.0176 respectively. Careful manual inspection of gene hits is therefore still necessary to identify novel hits associated with streptomycin-hypersensitivity.

### Identification of genes whose disruption conferred a fitness advantage under sub-inhibitory doses of streptomycin

We next focused on genes identified from our screen whose disruption conferred a fitness advantage, as indicated by the enrichment of transposon insertions (log_2_FC ≥ 2, q-value ≤ 0.05) in the streptomycin-treated library compared to the control in those genes (**Table 2**). Several of the top hits were members of the *nuo* operon, which encodes for the respiratory complex I. Loss-of-function mutations of this complex is believed to impair the proton gradient required for aminoglycoside uptake at the inner membrane, thus promoting aminoglycoside tolerance but not resistance (12, 36, 57). Another hit was *cyoE*, which encodes a subunit to the cytochrome b_0_ oxidoreductase (58), a terminal respiratory complex that generates PMF (59). Similar to respiratory complex I, disruption of cytochrome b_0_ oxidoreductase is expected to impair proton gradient generation and thus confer aminoglycoside resistance (60, 61). We also identified *arcA*, which belongs to the Arc two-component system that is involved in the regulation of respiratory complexes, where its disruption was indeed previously shown to promote aminoglycoside resistance (11). Another hit we found was the ribosome methyltransferase *rsmG*, whose mutants were previously associated with streptomycin resistance (62). One novel hit not previously implicated with aminoglycoside tolerance or resistance was *yebY*, a component of the YobA-YebY-YebZ copper uptake system (63). Interestingly, loss of this system results in the downregulation of the *mar* operon and subsequently increased sensitivity towards various antibiotics (but aminoglycosides were not tested) (64). Another novel hit was the flavin reductase *fre* (65), whose enzyme has been implicated as a source for superoxide, a reactive oxygen species (66). Loss of *fre* should result in lower intracellular concentrations of superoxide and subsequently decreased susceptibility against aminoglycosides (10, 12), although experimental evidence with *fre* mutants are lacking. Among the other hits were either small genes or genes essential in our LB-control library, including *iraM*, *obgE*, *pnp* and *yrfF* (now *igaA*). Manual inspection of these genes showed sparse insertions between the streptomycin-treated and control libraries.

**Table 2.** Genes identified to confer fitness advantage (i.e. resistant or tolerant; log_2_FC. ≥ **2 and q-value** ≤ **0.05) towards streptomycin when disrupted, with their BioTraDIS-predicted essentiality status in the control TIS experiment.**

| Gene | Function | $\log_2FC$ | logCPM | q-value | Ess. in control? |
| --- | --- | --- | --- | --- | --- |
| *argW* | arginine-tRNA synthetase | 6.32 | 1.81 | 4.35E-04 | N |
| *ackA* | acetate kinase A and propionate kinase 2 | 6.09 | 1.67 | 1.23E-03 | Y |
| *yoel* | uncharacterized protein | 5.07 | 2.63 | 4.42E-04 | Ambi. |
| *iraM* | RpoS stabilizer during Mg starvation, anti-RssB factor | 4.13 | 1.88 | 2.77E-03 | Y |
| *arcA* | response regulator in two-component regulatory system with ArcB or CpxA | 3.39 | 3.17 | 4.56E-04 | N |
| *mgrB* | regulatory peptide for PhoPQ, feedback inhibition | 3.24 | 2.95 | 1.27E-03 | N |
| *yebY* | DUF2511 family protein | 3.16 | 4.15 | 2.20E-06 | N |
| *nuoK* | NADH:ubiquinone oxidoreductase, membrane subunit K | 3.11 | 3.81 | 3.73E-04 | N |
| *obgE* | GTPase involved in cell partitioning and DNA repair | 3.10 | 2.43 | 1.72E-03 | Y |
| *nuoE* | NADH:ubiquinone oxidoreductase, chain E | 2.98 | 4.63 | 5.58E-05 | N |
| *nuoC* | NADH:ubiquinone oxidoreductase, fused CD subunit | 2.72 | 7.11 | 9.83E-17 | N |
| *argY* | arginine-tRNA synthetase | 2.55 | 3.82 | 7.22E-05 | N |
| *pnp* | polynucleotide phosphorylase/polyadenylase | 2.45 | 6.10 | 9.69E-09 | Y |
| *yrfF* | inner membrane protein | 2.45 | 3.59 | 2.74E-03 | Y |
| *nuoG* | NADH:ubiquinone oxidoreductase, chain G | 2.40 | 8.14 | 1.36E-16 | N |
| *nuoF* | NADH:ubiquinone oxidoreductase, chain F | 2.38 | 6.62 | 2.31E-10 | N |
| *nuoB* | NADH:ubiquinone oxidoreductase, chain B | 2.32 | 5.97 | 5.50E-07 | N |
| *rsmG* | 16S rRNA m(7)G527 methyltransferase, SAM-dependent; glucose-inhibited cell-division protein | 2.29 | 10.78 | 8.87E-28 | N |
| *yciS* | DUF1049 family inner membrane protein, function unknown | 2.25 | 2.71 | 4.03E-03 | N |
| *cyoE* | protoheme IX farnesyltransferase | 2.15 | 8.13 | 1.63E-21 | N |
| *fre* | NAD(P)H-flavin reductase | 2.15 | 8.28 | 6.22E-24 | N |
| *pta* | phosphate acetyltransferase | 2.06 | 4.29 | 1.41E-03 | N |
| *nuoN* | NADH:ubiquinone oxidoreductase, membrane subunit N | 2.01 | 6.91 | 1.29E-11 | N |

### Streptomycin sensitivity validation of selected hits from TIS screen

To validate our identified streptomycin-conditionally essential genes, we exposed single gene deletion mutants from the Keio collection to a sub-inhibitory concentration of streptomycin (35). Overnight cultures of the respective mutants (and parent wildtype strain; WT) were 10-fold serially diluted and inoculated onto LB agar with or without a sub-inhibitory concentration of streptomycin. Most of these mutants did indeed display strong streptomycin sensitivity compared to the wildtype, with a 10^5^-fold decrease in colony forming units (CFUs) (**Figure 2C**). Previously validated genes, such as *atpA*, *atpD*, *secG*, and *hflC* were indeed shown to be Strep sensitive (4, 11, 28). Additionally, we confirmed streptomycin sensitivity in mutants of novel genes discovered from our screen, including *yggT* and *yciB*. The *envC*, *cpxR* and *trkA* mutants only displayed modest reduction in CFU (*envC*: 10-fold, *cpxR*: 10-fold, or *trkA*: 10^2^-fold respectively) in the presence of streptomycin. The *waaC* and *qseC* mutants did not display streptomycin sensitivity. A possible reason could be the presence of secondary mutations in these Keio mutants, as it has been noted in other mutants in the Keio collection (19), such that they suppress the streptomycin sensitivity phenotype.

We also exposed selected mutants of genes whose disruption resulted in low level resistance/tolerance found in our screen. We found that mutants of *arcA*, *yebY*, *fre* and *rsmG* but not *nuoK* were less sensitive to streptomycin than the WT, with at least 10,000-fold increase in CFUs on LB supplemented with 20 µg/ml streptomycin, relative to the WT (**Figure 3**).

**Figure 3.**
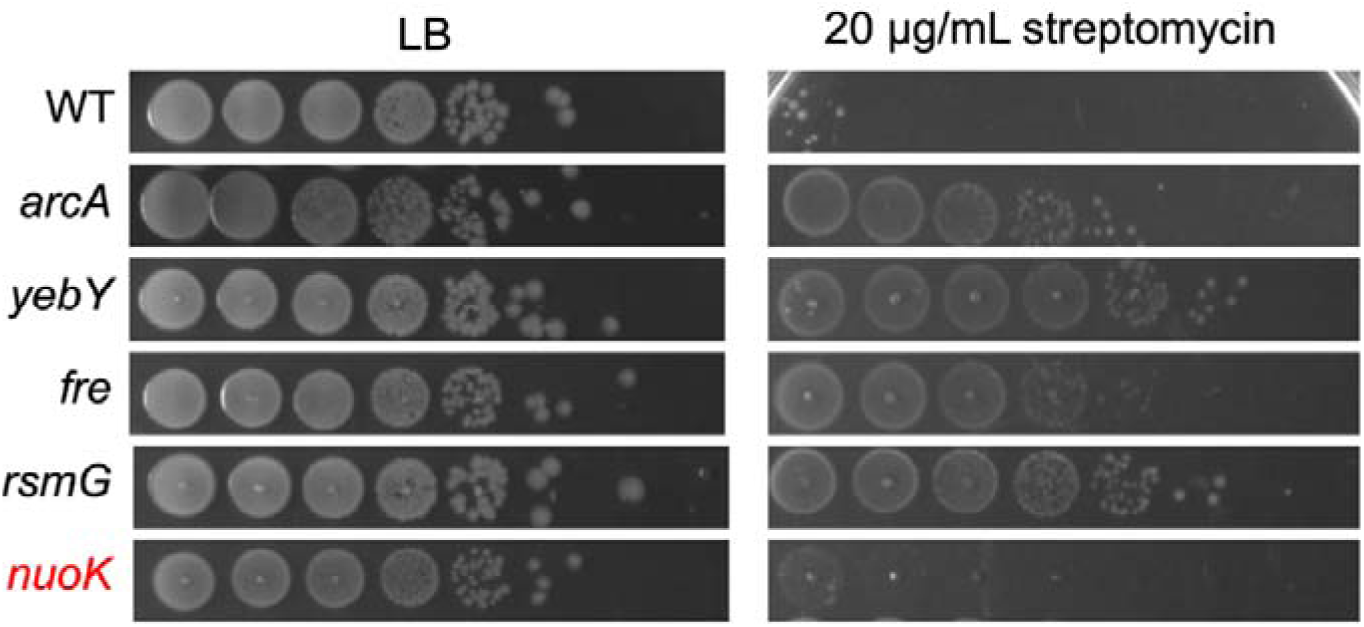
Validation of gene hits whose disruption confers low-level resistance towards streptomycin. Overnight cultures were normalised to OD_600_ = 0.1, ten-fold serially diluted, before being inoculated onto LB agar supplemented with 20 µg/ml streptomycin. Mutants that were not sensitive to streptomycin are highlighted in red.

### Streptomycin sensitive mutants generally do not display a hyperpolarised IM as indicated by membrane potential dye experiments

We observed that many of the streptomycin conditionally-essential genes identified in our screen were functionally linked to maintenance of the PMF, which is required for uptake of aminoglycosides across the IM (7). Given the central role of PMF in energising aminoglycoside transport, we hypothesised that the susceptible mutants possess a hyperpolarised IM, thereby enhancing streptomycin uptake and in turn increasing sensitivity to streptomycin. To test this, we used the fluorescent membrane potential dye DiOC_2_(3) to assess the relative membrane potential of the tested mutants (67). In the presence of a polarised IM (more positively charged on the outer leaflet relative to inner leaflet), the cationic DiOC_2_(3) traverses the IM and accumulates in the cytoplasm, causing a redshift of its emission wavelength (68), while, a higher-than-WT redshifted emission intensity indicates IM hyperpolarisation (67). As a depolarised control, WT cells were treated with the protonophore CCCP, where we saw a significant drop in redshifted emission intensity relative to untreated WT over time (**Figure 2B**). Across the WT and mutants, the redshifted emission intensities gradually decreased over 60 min, indicative of their membrane potential becoming depolarised presumably due to toxicities arising from prolonged DiOC_2_(3) exposure. Deletions of *yciB*, *yfgM*, *yggT* and *ppiD* did not significantly alter membrane potential relative to WT cells, indicating that their susceptibility to aminoglycosides is likely independent of membrane potential perturbations. Throughout the assay duration, the *envC* mutant was the only strain that exhibited higher emission intensities than WT. Interestingly, the *trkA* mutant possesses initially lower emission intensities than WT cells, in line with a previous study (69), but later plateaued at measurements higher than WT. Deletion of the Sec translocon components (*yajC*, *secG*), and *sdaC* resulted in consistently lower redshifted emission intensities which suggests that they possess more depolarised membranes relative to the WT. Surprisingly, we found similar results for the ATP synthase mutants (*atpA* and *atpD*), which contradict the prevailing hypothesis that these mutants possess hyperpolarised membranes to facilitate increased uptake of aminoglycosides (70). Taken together, this suggests that *E. coli* possess a myriad of mechanisms to survive streptomycin exposure that are downstream of the PMF-dependent uptake step.

### Overexpression of the IM protein SbmA confers hypersensitivity to streptomycin

It is not well understood why the PMF is required for aminoglycoside IM uptake. One hypothesis is that aminoglycosides hijack a PMF-dependent IM transporter for uptake (4). We hypothesise that overexpression of such a transporter would result in hypersensitivity towards aminoglycosides. The mini-Tn*5* used in our *E. coli* BW25113 TIS library possesses transcriptional readthrough activity that originates from the strong constitutive promoter of its chloramphenicol acyltransferase (*cat*) resistance cassette (30). Thus, for a given gene, any upstream insertion where the transposon (and *cat* promoter) is sense-oriented to its coding sequence should result in overexpression of that gene.

We compared sense insertions 100 bp upstream of every gene between our streptomycin treated and control libraries using the AlbaTraDIS package (71) (**Table 3**). We found 20 genes whose upstream sense insertions were depleted (log_2_FC ≤ −2, q-value ≤ 0.05). On manual inspection of each of these genes, we found that most had an immediate upstream gene that had sparse transposon insertions in both sense and antisense orientations. Particularly, some of these upstream genes include those identified earlier as streptomycin-susceptible, such as *atpF*, *cpxR*, *yggT* and *hflK*. Some of the upstream genes, such as *yidD*, *wecE* and *ygiB*, had more sparse insertions (in both sense and antisense orientations) in the streptomycin-treated library than the control, but were still deemed as non-essential in streptomycin-treated library. Thus, these hits were likely not the result of their overexpression *per se* but rather the disruption of their upstream gene partners conferring streptomycin susceptibility. For the remaining gene hits that did not have an upstream gene partner, we focussed on those encoding for IM proteins, which were *yqjA*, *ybgM* and *sbmA*. YqjA is part of the DedA family and may possess undecaprenyl phosphate flippase activity in a PMF-dependent fashion (72, 73). Given the structural dissimilarity between undecaprenyl phosphate and aminoglycosides, we concluded that *yqjA* was not a likely candidate and decided to not further validate it. YbgM is an uncharacterised IM protein, although proteomic studies have failed to detect its presence (74). Lastly, SbmA is an IM transporter protein with high substrate promiscuity, being able to uptake various antimicrobial peptides in a PMF-dependent fashion (75–79) (**Figure 4B**). Given that *sbmA* was our strongest hit (log_2_FC = −9.5, q-value = 4.41 x 10^−8^) with a clearly visible decrease in insertions upstream of the CDS (**Figure 4A**), and its substrate promiscuity, we decided to further investigate.

**Figure 4.**
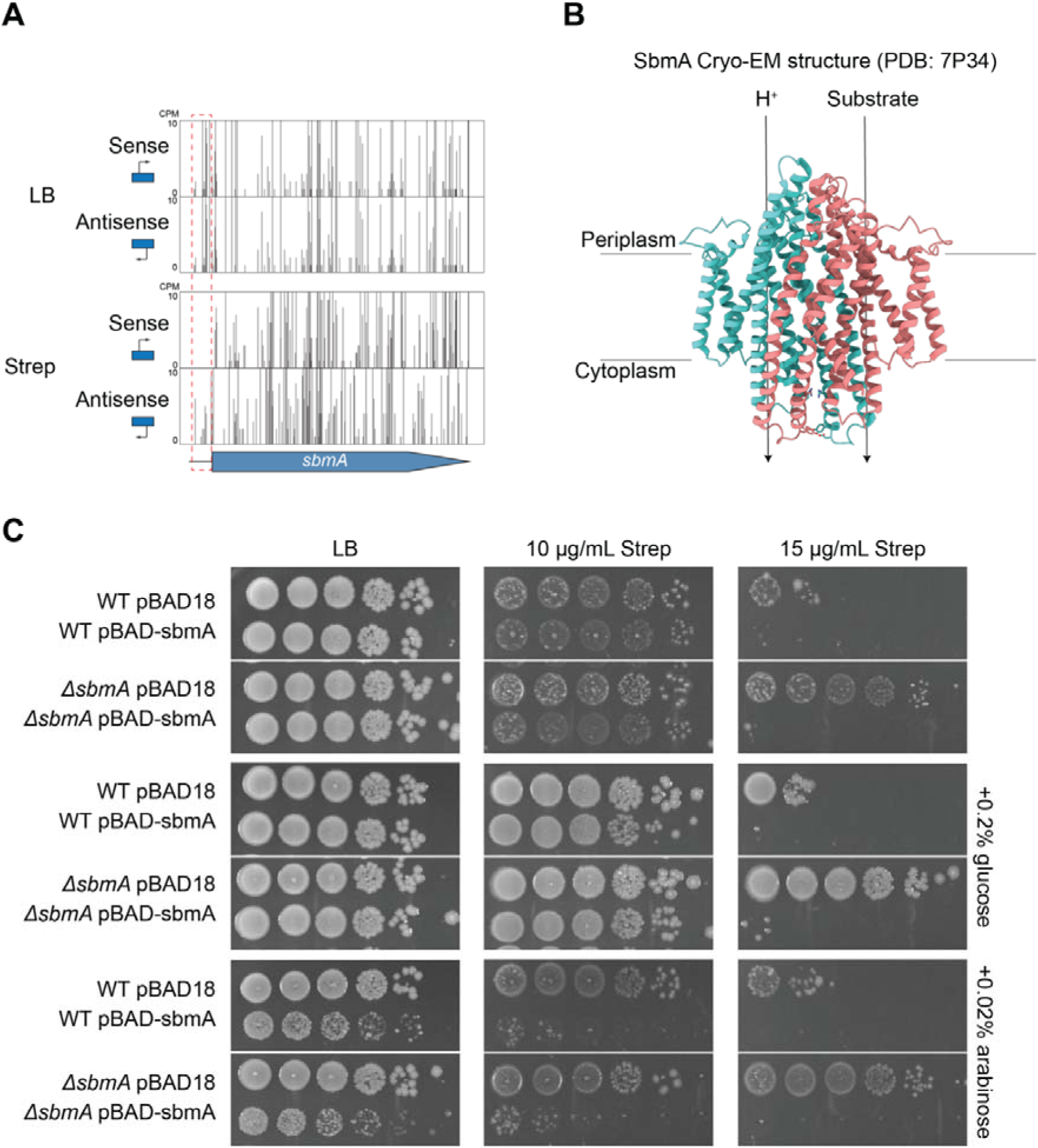
Ectopic overexpression of *sbmA* results in increased streptomycin sensitivity. **(A)** TIS plots separated based on the direction of the transposon’s antibiotic cassette promoter to *sbmA* – sense insertions correlate with the antibiotic cassette promoter is in the same orientation to *sbmA* and vice versa for antisense insertions. For clarity, a red-dashed box (spanning 100 bp) was added to indicate region where sense insertions were depleted in the streptomycin-treated library. **(B)** Cryo-EM structure (PDB: 7P34) of *E. coli* SbmA, a homodimeric (monomers in green and pink) IM transporter implicated for importing substrates such as antimicrobial peptides in a PMF-dependent fashion. **(C)** *E. coli* BW25113 WT or Δ*sbmA* deletion mutants transformed with a plasmid encoding for an arabinose-inducible copy of *sbmA* (pBAD-sbmA). Overnight cultures of the respective strains were diluted to OD_600_ = 0.1, ten-fold serially diluted and inoculated onto LB agar + carbenicillin plates supplemented with different concentrations of streptomycin, 0.02% arabinose or 0.2% glucose.

**Table 3.** Genes with a significant depletion or enrichment of insertions upstream of the CDS. Only insertions that were oriented in the sense direction relative to the to the genes’ respective coding sequences were tallied.

| Gene | Function | log<sub>2</sub>FC | logCPM | q-value | Upstream gene |
| --- | --- | --- | --- | --- | --- |
| *sbmA* | microcin B17 transporter | -9.57 | 4.17 | 4.41E-08 | - |
| *atpH* | F1 sector of membrane-bound ATPsynthase and delta subunit | -9.50 | 4.08 | 1.53E-07 | *atpF* |
| *cpxA* | sensory histidine kinase in two-component regulatory system with CpxR | -9.12 | 3.72 | 9.24E-06 | *cpxR* |
| *iclR* | transcriptional repressor | -8.23 | 2.89 | 1.41E-02 | - |
| *yggU* | UPF0235 family protein | -4.94 | 4.92 | 2.75E-08 | *yggT* |
| *yidC* | membrane protein insertase | -4.86 | 3.91 | 2.08E-03 | *yidD* |
| *malM* | maltose regulon periplasmic protein | -4.85 | 3.62 | 8.94E-03 | - |
| *atpA* | F1 sector of membrane-bound ATP synthase and alpha subunit | -4.78 | 3.81 | 4.16E-03 | *atpH* |
| *hflC* | modulator for HflB protease specific for phage lambda cII repressor | -4.50 | 4.97 | 7.60E-09 | *hflK* |
| *wzxE* | O-antigen translocase | -4.07 | 3.64 | 1.92E-02 | *wecE* |
| *ygiC* | ATP-Grasp family ATPase | -3.44 | 4.36 | 2.67E-03 | *ygiB* |
| *mlaD* | ABC transporter maintaining OM lipid asymmetry and anchored periplasmic binding protein | -3.21 | 4.27 | 3.72E-03 | *mlaE* |
| *yqjA* | general envelope maintenance protein; DedA family inner membrane protein | -3.12 | 4.40 | 3.02E-02 | - |
| *sthA* | pyridine nucleotide transhydrogenase and soluble | -3.10 | 4.64 | 3.06E-03 | - |
| *hslU* | molecular chaperone and ATPase component of HslUV protease | -2.87 | 4.44 | 3.82E-03 | *hslV* |
| *lrp* | leucine-responsive global transcriptional regulator | -2.80 | 4.14 | 4.64E-02 | - |
| *wzzE* | Entobacterial Common Antigen (ECA) polysaccharide chain length modulation protein | -2.76 | 4.16 | 3.38E-02 | *wecA* |
| *ybjM* | inner membrane protein | -2.42 | 5.05 | 2.45E-02 | - |
| *ampD* | 1 and 6-anhydro-N-acetylmuramyl-L-alanine amidase and Zn-dependent; murein amidase | -2.30 | 4.55 | 4.03E-02 | - |
| *waaL* | O-antigen ligase | -1.99 | 5.63 | 3.72E-02 | *wecC* |

To confirm that *sbmA* overexpression confers a streptomycin hypersensitivity phenotype, we cloned *sbmA* into the pBAD18 vector (80), termed pBAD-*sbmA*, where it is regulated under the arabinose-inducible P_BAD_ promoter. Thus, the supplementation of arabinose into the media would result in the induction of *sbmA*. Conversely, supplementation with glucose results in the repression of plasmid-based *sbmA*. Overnight cultures of *E. coli* BW25113 WT cells harbouring pBAD-*sbmA* or control plasmid pBAD18 were then serially diluted and inoculated onto LB agar containing the various supplements. We observed that *E. coli* pBAD-*sbmA* colonies appeared smaller relative to strains harbouring the empty pBAD vector when the LB medium was supplemented with arabinose (**Figure 4C**). This might suggest overexpression of *sbmA* is toxic, however, no significant decrease in CFUs was observed from overexpressing *sbmA*.

Next, we exposed *E. coli* pBAD-sbmA strains to sub-inhibitory concentrations (10 μg/mL) of streptomycin. In the presence of 10 μg/mL streptomycin and arabinose, we observed a 10^4^-fold reduction of *E. coli* pBAD-sbmA CFUs compared to the *E. coli* pBAD18 control. Conversely, no changes of CFUs were seen when growing the same strains on 10 μg/mL streptomycin and glucose repression (**Figure 4C**). These results support the notion that SbmA overexpression does indeed confer streptomycin hypersensitivity. We then tested whether the low-level streptomycin resistance phenotype found in the Δ*sbmA* mutant could be complemented by introducing pBAD-sbmA (**Figure 4C**). However, ectopic overexpression of *sbmA* in Δ*sbmA* cells were sensitive to both sub-inhibitory concentrations (10 µg/mL) and inhibitory (15 μg/mL) concentrations of streptomycin. Furthermore, glucose-mediated repression still rendered Δ*sbmA* cells sensitive towards inhibitory doses (15 µg/ml) of streptomycin than WT-levels (i.e. WT pBAD18). This suggests that the residual *sbmA* expression levels arising from the pBAD-*sbmA* vector exceeds *sbmA*’s native expression levels.

### Mutants overexpressing sbmA do not have an envelope defect

We noticed that when plasmid-encoded *sbmA* was overexpressed via arabinose induction, its colonies appeared to be smaller than WT sizes (**Figure 4C**). This defect could be indicative of membrane permeability defects and could thus explain their streptomycin hypersensitivity phenotypes. To explore this possibility, we tested the susceptibility of these strains towards vancomycin. Vancomycin exerts its antimicrobial activity by binding to the peptide crosslinks in the peptidoglycan layer, but cannot traverse through an intact OM. As such, only cells with a compromised cell envelope would be sensitive towards vancomycin and subsequently a reduction in colony numbers should be observed. In the presence of vancomycin but not arabinose, the colony sizes of all strains were smaller than when untreated (**S. Fig. 3**). With the combined presence of vancomycin and ectopic *sbmA* overexpression, colonies were translucent and even smaller, although neither the WT nor Δ*sbmA* background had a significant reduction in colony numbers. Thus, we conclude that the streptomycin sensitivity conferred from *sbmA* overexpression was unlikely attributable to membrane permeability defects, and the colony shrinkage phenotype from *sbmA* overexpression could simply be indication of growth inhibition, as noted before (81).

### Loss of SbmA confers decreased susceptibility to aminoglycosides, but not other antibiotics

Loss of *sbmA* was previously shown to confer resistance to the antimicrobial peptides it is implicated for importing (75). We hypothesised this might also apply to aminoglycosides. In our TIS data, we only noticed a modest but statistically significant enrichment of insertions within the *sbmA* coding sequence (log_2_FC = 1.3, q-value = 9.00 x 10^−11^) in the streptomycin-treated library. To validate this, we exposed the BW25113Δ*sbmA* mutant (Δ*sbmA*) to an inhibitory concentration of streptomycin as well as other aminoglycosides including kanamycin, gentamicin and tobramycin (**Figure 4**). Indeed, Δ*sbmA* grew better than its parent WT strain. However, the Δ*sbmA* mutant was sensitive to the same aminoglycosides at higher concentrations, suggesting the presence of additional proteins or mechanisms besides SbmA to import aminoglycosides in a PMF-dependent fashion.

Given the substrate promiscuity of SbmA for hydrophilic compounds, SbmA could also promote the uptake of other similar-sized antibiotics and decreased susceptibility in these antibiotics should be observed in cells lacking *sbmA*. We exposed the Δ*sbmA* mutant against inhibitory doses of chloramphenicol, ampicillin and tetracycline but no resistance phenotype to those antibiotics was observed (**Figure 5**). As Δ*sbmA* retains WT-level sensitivities to chloramphenicol and tetracycline, both of which target the ribosome, its observed decreased sensitivity only to aminoglycosides suggests that loss of *sbmA* is unlikely to simply promote survival through a mechanism downstream of ribosomal stress. Taken altogether, our findings indicate that SbmA facilitates the IM uptake of aminoglycosides rather than functioning as a generalised importer for diverse antimicrobial compounds.

**Figure 5.**
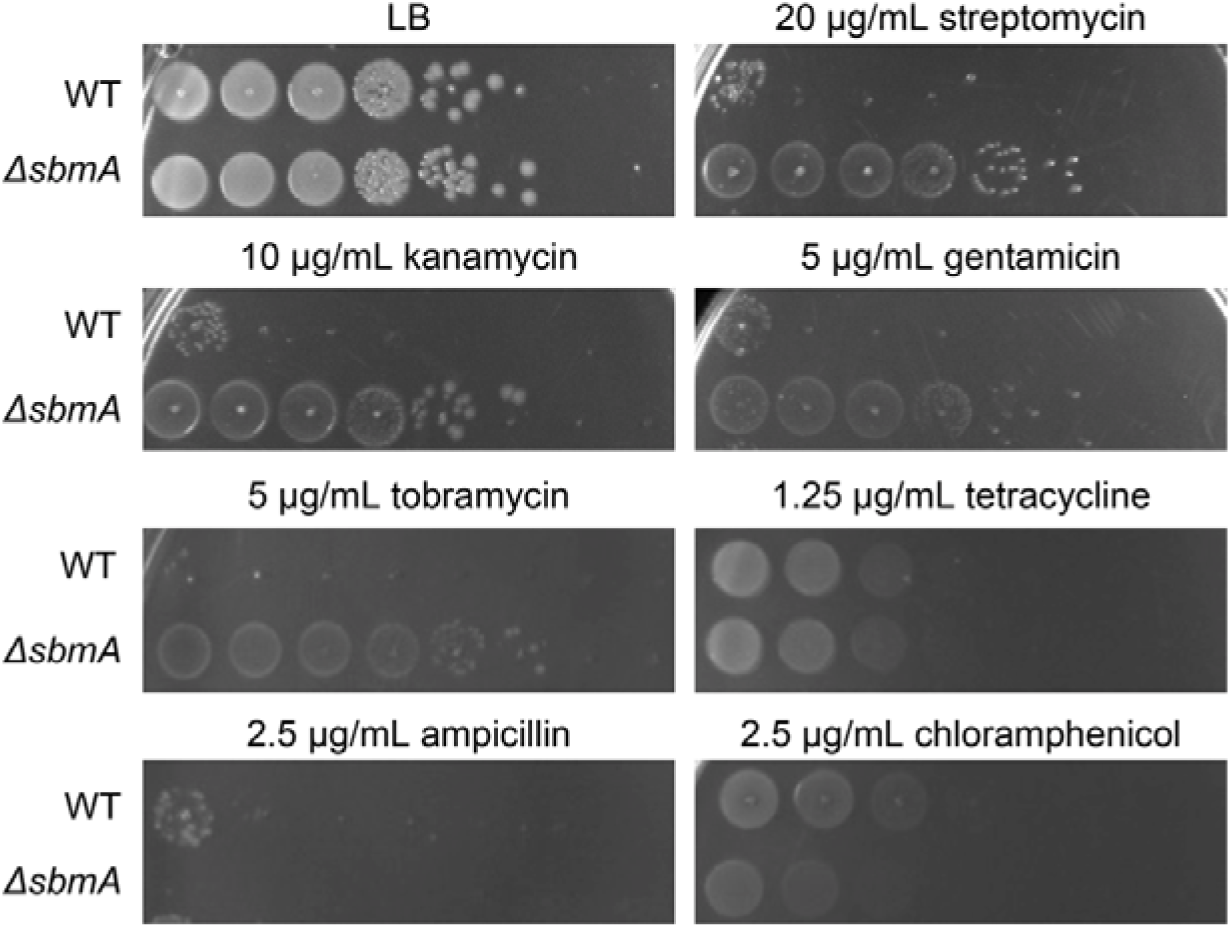
Loss of *sbmA* confers low level resistance against different aminoglycosides but not tetracyclines, β-lactams and chloramphenicol. Overnight cultures of *E. coli* BW25113 WT or its Δ*sbmA* deletion mutant were diluted to OD_600_ = 0.1, ten-fold serially diluted and spotted onto LB agar supplemented with inhibitory concentrations of streptomycin, kanamycin, gentamicin, tobramycin, tetracycline, ampicillin and chloramphenicol.

## Discussion

In this work, we have used TIS to uncover the genetic basis of the streptomycin susceptibility and low-level resistance in model *E. coli*. Demonstrating the robustness of our TIS screen, we have identified links between streptomycin action and known genes that confer streptomycin hypersensitivity when disrupted, including the ATP synthase, cell division and protein translocation pathways across the inner membrane. Additionally, we uncovered novel links between several inner membrane transporter proteins, including *sdaC* and *trkA*, as well as uncharacterised genes such as *yciB* and *yggT*. We also found several genes that gave rise to low levels of streptomycin resistance or tolerance when disrupted, including components of the respiratory complex I (*nuo*) and the Arc two-component system, many of which were previously reported (11, 36, 57). However, our screen also identified novel hits such as *yebY* and *fre* that conferred streptomycin resistance or tolerance. Lastly, we identified an inner membrane transporter, SbmA, which when disrupted conferred some resistance towards several aminoglycosides but not other ribosomal-targeting antibiotics such as chloramphenicol and tetracycline, while its overexpression conferred streptomycin hypersensitivity.

It is largely unclear as to why gene disruption of members of the Sec translocon (e.g. SecG and YajC) could confer hypersensitivity to streptomycin, as identified in our screen. It is believed that upon aminoglycoside-induced mistranslation, the Sec translocon embeds nascent mistranslated proteins into the IM, resulting in the formation of pores that cause further uptake of aminoglycosides (11). Furthermore, the Sec translocon also traffics numerous periplasmic proteases that degrade misfolded proteins, including PpiD which were also found as conditionally essential in our screen. Thus, disruption of Sec activity would expectedly hamper the export of these proteases to aid in reducing the abundance of these mistranslated protein pores. Furthermore, Sec activity is also dependent on PMF (in addition to ATP) for proper trafficking of client proteins (82, 83), although the link between such dependence and aminoglycoside uptake is poorly explored.

In our screen, we also surprisingly identified cell division-associated genes linked to increased sensitivity to streptomycin, including EnvC, MinCDE and YciB. EnvC is a periplasmic factor that localises at mid-cell and activates amidase (AmiA and AmiB) activity essential for cytokinesis. Loss of *envC* results in cell filamentation and envelope defects (84). Thus, we hypothesise filamentation from *envC* disruption allows for a greater surface area accessible for streptomycin to enter the cell, alongside with a compromised cell envelope, which results in increased sensitivity towards streptomycin. MinCDE regulates proper divisome formation at the midcell by colocalising in cell poles and preventing FtsZ filaments to form there. It was previously shown that MinD requires PMF (in particular, the transmembrane potential component) for correct colocalisation in *Bacillus subtilis* (85). Should such findings be generalisable to *E. coli*, a simple explanation to link streptomycin hypersensitivity and loss of MinCDE would be that such cells would have ‘unconsumed’ PMF and the hyperpolarised membrane would subsequently allow for increased streptomycin uptake. Finally, the poorly characterised *yciB* was found to interact with several key cell division and elongation-associated proteins including RodZ, MurG and MreC (44), although to what extent does the loss of *yciB* would affect the function of these proteins remains unknown.

PMF is intricately linked to aerobic respiration and thus any perturbations to the latter should affect aminoglycoside activity (65). Indeed, loss-of-function mutations in components of ATP synthase, also identified in our TIS screen, results in hypersensitivity to streptomycin and other aminoglycosides (4, 7, 18, 86). It is believed that the loss of ATP synthase activity results in the uncoupling of oxidative phosphorylation and PMF, resulting in a hyperpolarised membrane (87). Conversely, we also identified disruptions in the *nuo* operon, which encodes subunits of respiratory complex I, conferred low-level streptomycin tolerance/resistance. The loss of proton pumping activity in *nuo* mutants is believed to reduce the transmembrane potential (88), thereby limiting the uptake of aminoglycosides (12, 86). In support of this, adaptive laboratory evolution experiments in *E. coli* with various aminoglycosides (including streptomycin) yielded resistant mutants related to these respiratory complexes (89).

A simple explanation for the functionally diverse set of conditionally essential genes found in our screen would be that their disruption results in membrane hyperpolarisation, thereby facilitating greater streptomycin uptake. However, we did not observe any trend in membrane polarisation in various deletion mutants of these genes using the voltage-sensitive DiOC_2_(3) dye (68). In the staining protocol we used, we pre-incubated cells with EDTA to promote intake of DiOC_2_(3) by destabilising the LPS layer, a common practice done previously (67, 90, 91). Such destabilisation caused by EDTA could in turn activate stress responses, including the Cpx, Rcs (regulation of capsule biosynthesis) and Bae (bacterial adaptive response) pathways (92). Within these pathways, the Cpx regulon constitutes the expression of several respiratory complexes that could alter membrane potential (87, 93, 94), which could be exacerbated in different genetic backgrounds such as the knockout mutants we tested. Given the pleiotropic consequences of chelators, genetically-encoded sensors could serve as an attractive alternative, such as calcium-sensitive fluorescent proteins that could indirectly measure PMF (86, 95).

A longstanding question in understanding aminoglycoside mechanism of action is the role of PMF in aiding uptake (and by extension, bactericidal activity) of such antibiotics. A possible explanation is the presence of a PMF-utilising inner membrane transporter. In our screen, we identified one such candidate, SbmA. This protein belongs to a larger SbmA/BacA family, which comprises conserved peptide transporters that mediate the uptake of diverse antimicrobial and host-derived peptides, thereby influencing antimicrobial susceptibility, resistance, and host-microbe interactions across bacterial phyla (96). In *Enterobacteriaceae*, loss of SbmA confers resistance to many intracellularly acting peptide antibiotics (76, 78), whereas BacA homologues in alphaproteobacteria facilitate symbiosis and virulence through interactions with host antimicrobial peptides (97, 98). BacA homologues in *M. tuberculosis* also was uptake of bleomycin, aminoglycosides, and other hydrophilic compounds (98, 99). Structural studies have revealed SbmA possesses an ABC exporter-like fold with a large hydrophilic substrate-binding cavity that enables broad substrate specificity and proton-coupled transport (75, 76, 79, 100). Here we show that overexpression of this gene, even at residual levels, conferred hypersensitivity towards streptomycin, suggesting a possible role for SbmA in aminoglycoside uptake. This is in support of a recent study, where introduction of a synthetic antimicrobial peptide resensitised a gentamicin-resistant strain of *E. coli*, which was found to be concomitant with elevated expression levels of *sbmA* via qRT-PCR (101). We also found that deletion of *sbmA* resulted in modest but not complete resistance towards streptomycin, kanamycin, gentamicin and tobramycin. In recent adaptive laboratory evolution studies, *sbmA* mutants were consistently obtained in amikacin-exposed (102) or gentamicin-exposed (103) *E. coli* biofilm populations but not in planktonic populations. Given our experiments primarily worked with planktonic cultures and that deletion of *sbmA* only gave rise to low-level resistance (i.e. two-fold), we hypothesise that *sbmA* is poorly expressed in planktonic cultures, in line with previous translational reporter assay studies (81). Expression of *sbmA* is positively regulated by the induction of the σ^E^ extracytoplasmic (104) and Cpx (105) stress responses, both of which are induced in *E. coli* biofilms (106). In contrast, we speculate that planktonic cells possess alternative PMF-dependent and/or PMF-independent mechanisms to uptake aminoglycoside at the inner membrane, including inner membrane mechanosensitive channels, amino acid transporters and/or sugar transporters (102, 107–111). Thus, we believe that SbmA may play a role in the mechanism-of-action of streptomycin (and aminoglycosides) in *E. coli*, especially in biofilms, a more clinically relevant context.

Our work here highlights the utility of whole genome screens such as TIS for uncovering (and validating) contributions from genes of unknown function (23, 27, 112, 113), inlcuding *yciB*, *yggT* and *yebY*. We foresee our data to be a useful starting point to better understand the physiological relevance of these genes.

In conclusion, although streptomycin, and by extension aminoglycosides, are one of the most well characterised classes of antibiotics, the molecular mechanisms governing aminoglycoside action and uptake remains to be fully characterised. Our TIS screen data presented here reaffirms previous discoveries and identifies novel association to pathways, providing greater clarity to our understanding of the aminoglycoside class. This could serve as a stepping stone to develop adjuvant or combinatorial therapies against the increasing prevalence of antimicrobial resistant infections.

## Materials and Methods

### Bacterial strains and Media

Strains were routinely cultured in Luria Bertani (LB) broth (1% (w/v) NaCl, 1% (w/v) tryptone and 0.5% (w/v) yeast extract) or LB agar media (1% (w/v) NaCl, 1% (w/v) tryptone, 0.5% (w/v) yeast extract and 1.5% agar (w/v)) at 37°C with aeration and shaking when necessary. When antibiotics for mutant or clone selection are used, the final concentration was 50 µg/mL carbenicillin (Astral Scientific, Australia). *E. coli* K-12 strain BW25113 is the parent strain for the Keio single-gene deletion mutant library (35), and has the following genotype: F-, Δ*(araD-araB)567*, Δ*lacZ4787*(::rrnB-3), λ*^−^*, *rph-1*, Δ*(rhaD-rhaB)568*, *hsdR514*.

### Construction of plasmid constructs harbouring sbmA

The pBAD18 vector is an arabinose-inducible expression vector; pBR322 *ori*, *araC*, P_BAD_, Amp^R^ (80). The BW25113 *sbmA* gene was amplified by PCR and introduced into pBAD18 via *Hind*III and *Xho*I sites; *Hind*III and *Xho*I sites corresponds to the 5’-end and 3’-end in *sbmA* respectively. Oligos used in this study are listed in Table 4.

**Table 4.**
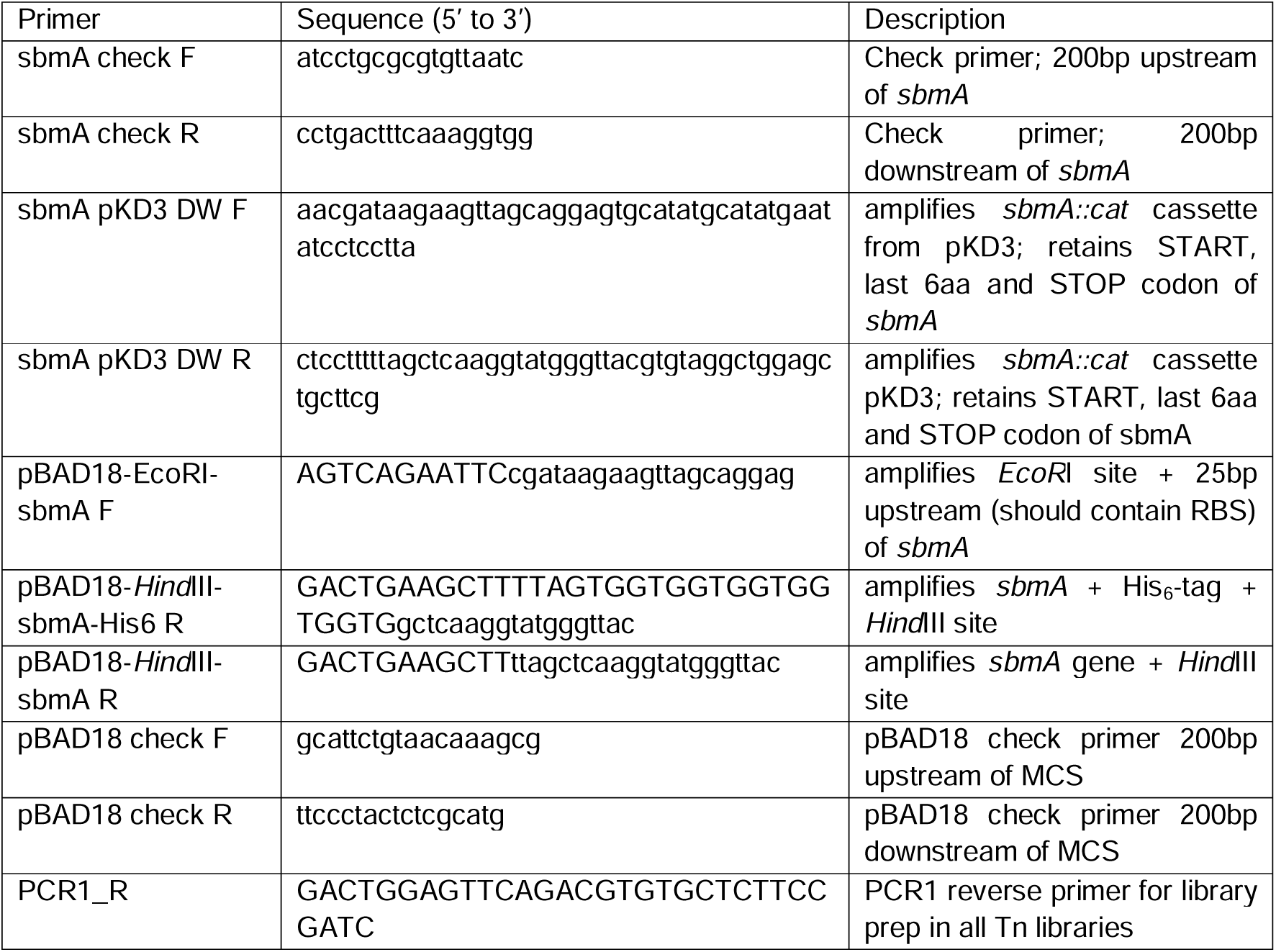
Oligos used in this study.

### Streptomycin MIC Assay

Overnight cultures, in biological triplicates, were normalised to a starting OD_600_ = 0.05, equivalent to ∼5 x 10^7^ colony forming units (CFU)/mL to enable sufficient coverage of the transposon mutants during the actual TIS library screen, in 50 mL LB media in 250 mL Erlenmeyer flasks. Flasks were then placed in a shaking incubator at 37 °C with shaking at 180 rpm, with OD_600_ measurements taken hourly using a UV-vis spectrophotometer.

### Streptomycin TIS screen

The *E. coli* BW25113 TIS library was constructed as previously described (30). To screen the *E. coli* BW25113 TIS library against sub-inhibitory doses of streptomycin, the library was inoculated into 50 mL LB media with or without 4 μg/mL streptomycin sulfate in 250 mL Erlenmeyer flasks to a starting OD_600_ of 0.05. Cultures for each condition were performed in duplicate. Cultures were incubated at 37 °C with shaking 180 rpm. OD_600_ measurements were periodically measured until an OD_600_ = 0.5 was reached (first passage). Next, 5 mL from each culture was transferred into 45 mL fresh LB containing the same supplements as it had and incubated with shaking until OD_600_ = 0.5 was reached (second passage). This passaging of cultures was performed once more (third passage). Cells were harvested when the cultures during the third passage reached an OD_600_ = 0.5 and were processed for genomic DNA extraction.

### Transposon sequencing

For each TIS library sample, two technical replicates were prepared for sequencing. Genomic DNA (gDNA) was extracted using the DNeasy Blood and Tissue kit (Qiagen, Germany) as per manufacturer instructions. gDNA was quantified, normalised to 1 µg in 130 μL nuclease free water and transferred into microTUBE-130 AFA (Adaptive Focused Acoustics) Fiber Screw-Cap tubes (Covaris, USA). Using a Covaris E220 sonicator (Covaris, USA), the gDNA was then sheared to approximately 300 bp fragments, using the following settings: peak power = 175, cycles/burst = 200, duty factor =100, duration (per sample) = 180 s. After sonication, samples were transferred into 1.5 mL tubes and concentrated to a volume of 50 μL using a vacuum concentrator. To enrich the transposon-gDNA junctions for Illumina high throughput sequencing, the subsequent steps use components of the NEBNext Ultra DNA Library Prep kit (E7645; New England Biolabs, USA) unless stated otherwise. First, fragmented DNA ends were repaired, and adaptor ligated. DNA fragments were then size-selected using SPRISelect magnetic beads (Beckman-Coulter, USA), a magnetic stand for bead-supernatant separation, and freshly prepared 80% (v/v) ethanol for washes (200 µL per wash). In this step, fragment size selection was double sided, with the fragments first mixed with beads in a 1:0.55 DNA:beads volume ratio. The supernatant was then transferred and mixed with fresh beads at a 6:1 supernatant:beads volume ratio to select for fragments sizing at 300-400 bp. The supernatant was discarded, and the beads were washed twice and eluted in 15 μL of Buffer EB (10 mM Tris pH 7.5). Transposon-gDNA junctions in the eluate were enriched via PCR using primers targeting the transposon’s 5′-end relative to the chloramphenicol cassette and the previously ligated adaptor. The following thermocycler conditions were used: 98 °C for 3 min, 10 cycles consisting of 98 °C for 15 s (denaturation), 65 °C for 30 s (annealing) and 72 °C for 30 s (extension), and a final extension of 72 °C for 1 min. Following another round of size selection (at a DNA:beads 0.9:1 volume ratio; discarding the supernatant and beads washed twice before eluted in 15 μL of Buffer EB), the PCR-enriched junctions were subjected to final round of PCR to introduce indices using NEBNext Multiplex Oligos for Illumina (New England Biolabs, USA) and a custom-designed barcoded primer. The custom primer features a 6 to 9 nucleotide barcode to introduce nucleotide complexity and staggered sequencing of the transposon’s first few nucleotides, as well as a 22 bp sequence homologous to the transposon 5′-end (TnChlor library). The following thermocycler conditions were used: 98 °C for 3 min, 20 cycles consisting of 98 °C for 15 s (denaturation), 65 °C for 30 s (annealing) and 72 °C for 30 s (extension), and a final extension of 72 °C for 1 min. The PCR product (now sequencing-ready libraries) was size selected again (at a DNA:beads 0.9:1 volume ratio; discarding the supernatant and beads washed twice before eluted in 35 μL of Buffer EB) and quantified by quantitative PCR (qPCR) (using thermal cycling conditions recommended by manufacturer) using the NEBNext Library Quant kit (New England Biolabs, USA) on the ViiA 7 Real-Time PCR System (Thermo Fisher Scientific, USA). The libraries were then diluted and denatured to a final concentration of 16 pM and spiked with 5% (v/v) of stock 20 pM PhiX library (Illumina, USA). Finally, fragments were sequenced on an Illumina MiSeq platform (Illumina., USA), using a MiSeq V3 150-cycle kit (Illumina Inc., USA).

### Processing of TIS reads to determine transposon insertion sites

Illumina reads were first trimmed of their 6 to 9 nucleotide barcode and subsequently transposon tags using the FastX toolkit (Coldspring Harbour Laboratories, USA). The Trimmomatic tool (114) was used to remove low quality (Phred score <33) and short (<20nt) reads. The remaining reads were then used as input into the bacteria_tradis pipeline from BioTraDIS to generate plot files depicting the position for every transposon insertion site in the library (115). In the pipeline, we used the SMALT algorithm to align reads to the *E. coli* BW25113 reference genome (NCBI GenBank accession: CP009273.1) using the following bacteria_tradis parameters: −mm 1 −m 0 −-smalt −-smalt_y 1 −-smalt_r 0. The output file contains the read depth (i.e. transposon insertion) for each bp of the *E. coli* BW25113 genome.

### Differential transposon insertion abundance analysis via edgeR

To compare the insertion abundance for each gene between TIS library samples, the tradis_comparison.R script in BioTraDIS was used with no minimal read filter set. The script uses the edgeR package (116) to compare normalised read counts (counts per million; CPM) between samples (each in technical duplicates), and therefore transposon mutant abundance alongside statistical significance. Statistical significance (p-value) were calculated via the binomial test and adjusted via the Benjamini-Hochberg procedure to derive q-values.

### Functional enrichment of TIS streptomycin screen hits

To identify genes required for streptomycin survival, we used the tradis_comparison.R script from BioTraDIS, and genes with log_2_FC < −2 (i.e. fourfold less normalised mapped reads in the treated sample relative to control) and q-value (q-value) < 0.05 were further used to perform Clusters of Orthologous Genes (COG) enrichment analysis on the eggNOG webserver (117).

### Validation of TIS streptomycin screen gene hits via testing of streptomycin susceptibility of knockout mutants

Overnight cultures of *E. coli* BW25113 or its gene-knockout derivatives in LB were first normalised to OD_600_ = 0.1 before 10-fold serial dilutions were made. The dilutions were then inoculated onto LB agar containing with or without antibiotics. Plates were incubated 37 °C for 16 h-18 h.

### Membrane potential assay

Overnight cultures were first sub-cultured 1:100 into 5 mL of LB media and grown until an OD_600_ of around 0.4. Cells were harvested by centrifuging 1 mL of cultures at 2400 x *g* for 10 min at room temperature, and each pellet was subsequently resuspended in 400 µL of phosphate-buffered saline (PBS; 137 mM NaCl, 2.7 mM KCl, 10 mM Na_2_HPO_4_, 1.8 mM KH_2_PO_4_, pH 7.4), which normalises the cell density to an OD_600_ = 1). To facilitate dye uptake, 8 µL 0.5 M EDTA (pH 8.0) was added to this suspension and incubated at room temperature for 5min before a second spin-down (2400 x *g* for 10min). The resulting pellet was then resuspended in 400 µL of LB supplemented with 30□μM DiOC_2_(3) (diluted from a stock solution of 3 mM DiOC_2_(3) in DMSO; powder obtained from Sigma-Aldrich, USA) 1 mM glucose and 1 mM CaCl_2_ to yield the final stained cell suspension. For measurement, 100 µL of each sample was loaded into the wells of a 96-well plate and allowed to incubate for at least 5 minutes at room temperature. As a depolarized control, 4 µL 500 mM carbonyl cyanide m-chlorophenyl hydrazone (CCCP) was additionally added to a separate 100 µL aliquot of stained cells. The membrane potential was then quantified using a plate reader by measuring the green (560 nm) and red (670 nm) fluorescence emission values from the same excitation wavelength of 490 nm. Results were expressed as a fraction of green fluorescence divided by red fluorescence. Lower fluorescence values indicate a lower (more negative) membrane potential.

## Acknowledgements

We thank the members of the Henderson lab for critical reading of the manuscript.

## Data Availability Statement

Sequencing data associated with this project would be available online.

---

**S. Table 1.** Sequencing reads metrics for TIS experiment samples.

|  | Reads<br>Mapped | Unique Insertion Sites |
| --- | --- | --- |
| Control Rep 1 | 1,934,869 | 365,068 |
| Control Rep 2 | 4,461,889 | 443,405 |
| **Combined** | **6,396,758** | **582,279** |
| streptomycin Rep 1 | 1,861,651 | 334,051 |
| streptomycin Rep 2 | 1,715,636 | 324,877 |
| **Combined** | **3,577,287** | **579,997** |

## Supplementary Figures

**Supplementary Figure 1.**
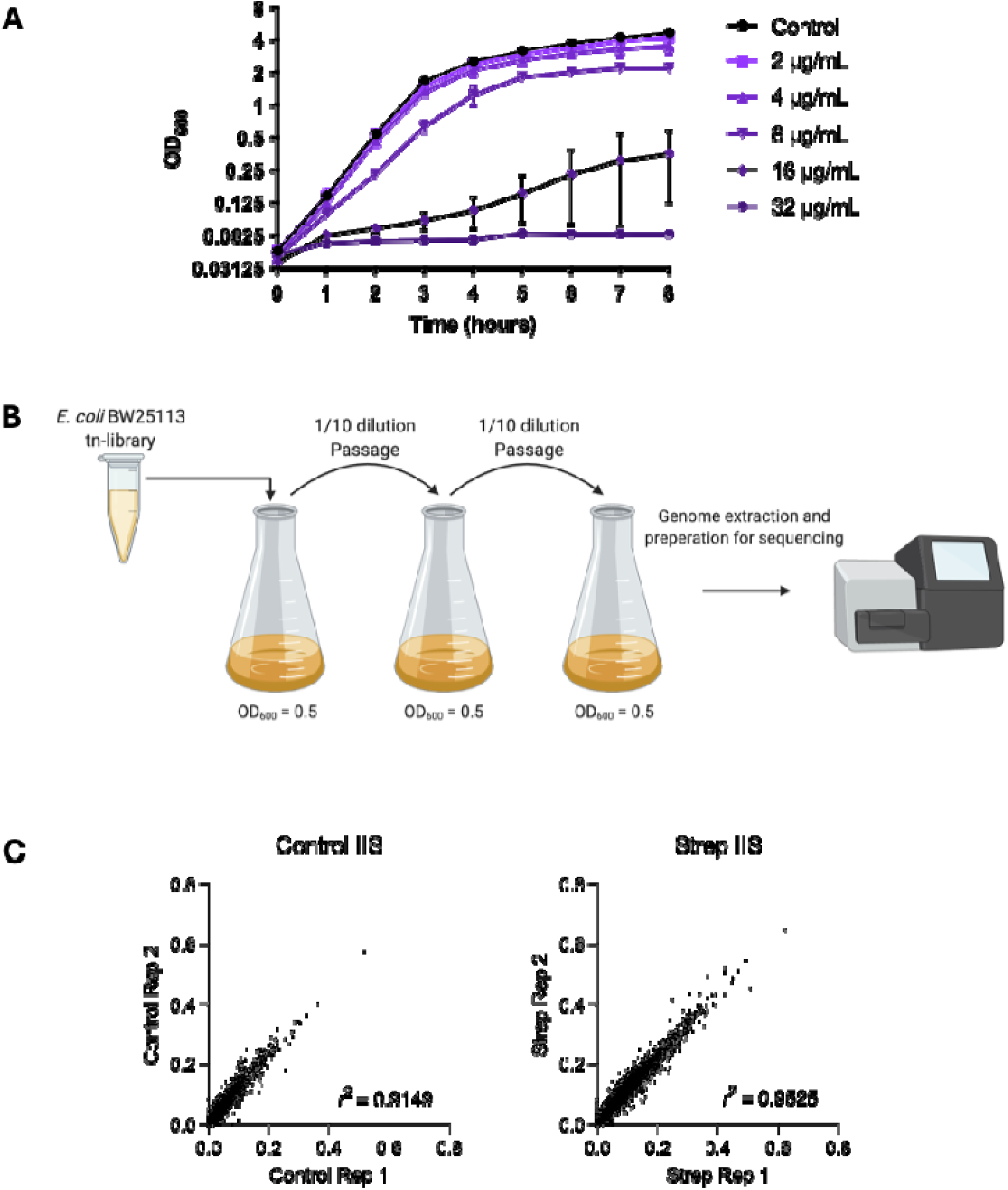
Growth conditions for transposon mutant streptomycin screen. **(A)** Growth curve displaying the optical density (OD_600_) of *E. coli* BW25113 grown in LB, in duplicate with or without streptomycin at varying concentrations. **(B)** Schematic of the experimental design for exposing the *E. coli* BW25113 transposon library to streptomycin. The library was grown in LB with or without 4 µg/ml streptomycin in duplicate for 3 passages. **(C)** Each point on the plot represents a gene and its insertion index score (the number of unique insertions within a gene divided by gene length) between each replicate per condition. Pearson correlation coefficient (*r*^2^) is indicated.

**Supplementary Figure 2.**
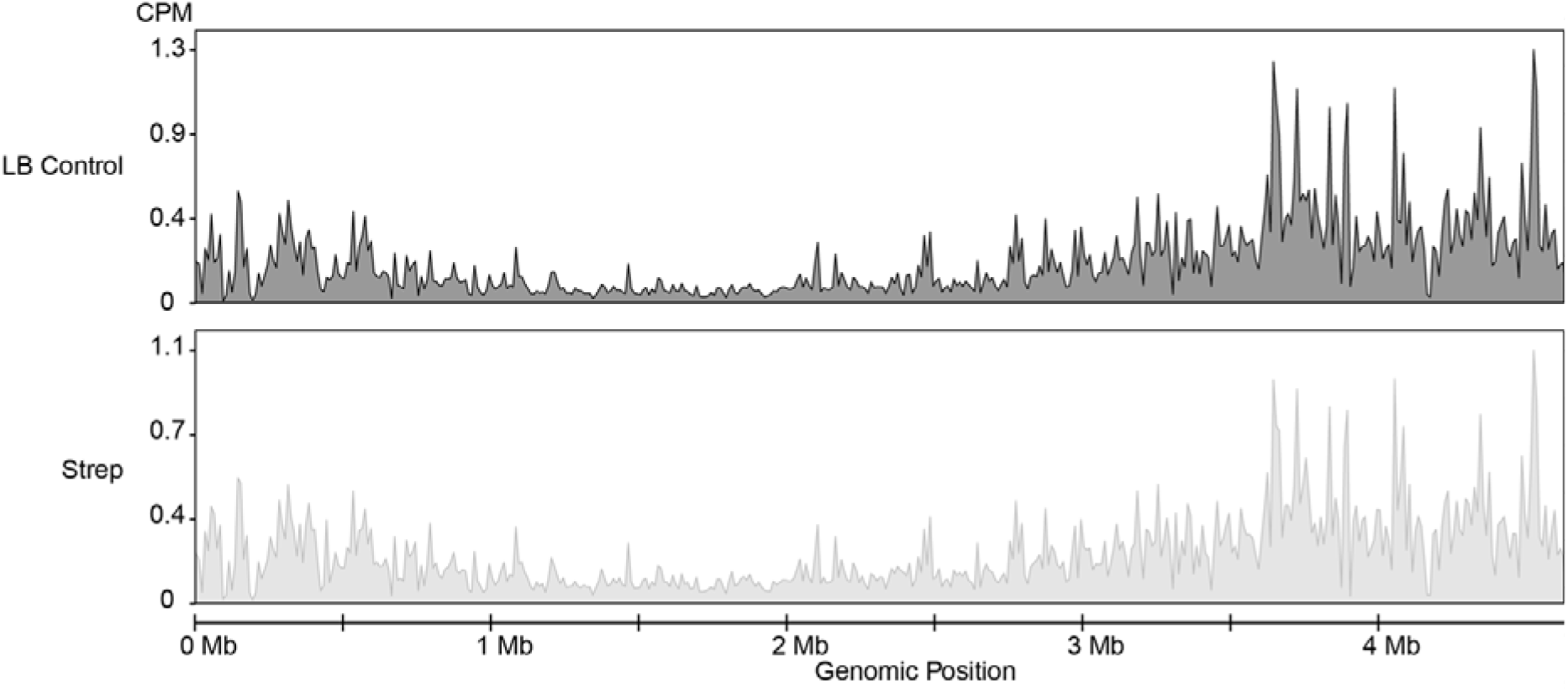
Sinusoidal plot of TIS data. Normalised insertions were binned into 10 kb windows and their density across the genome were visualised.

**Supplementary Figure 3.**
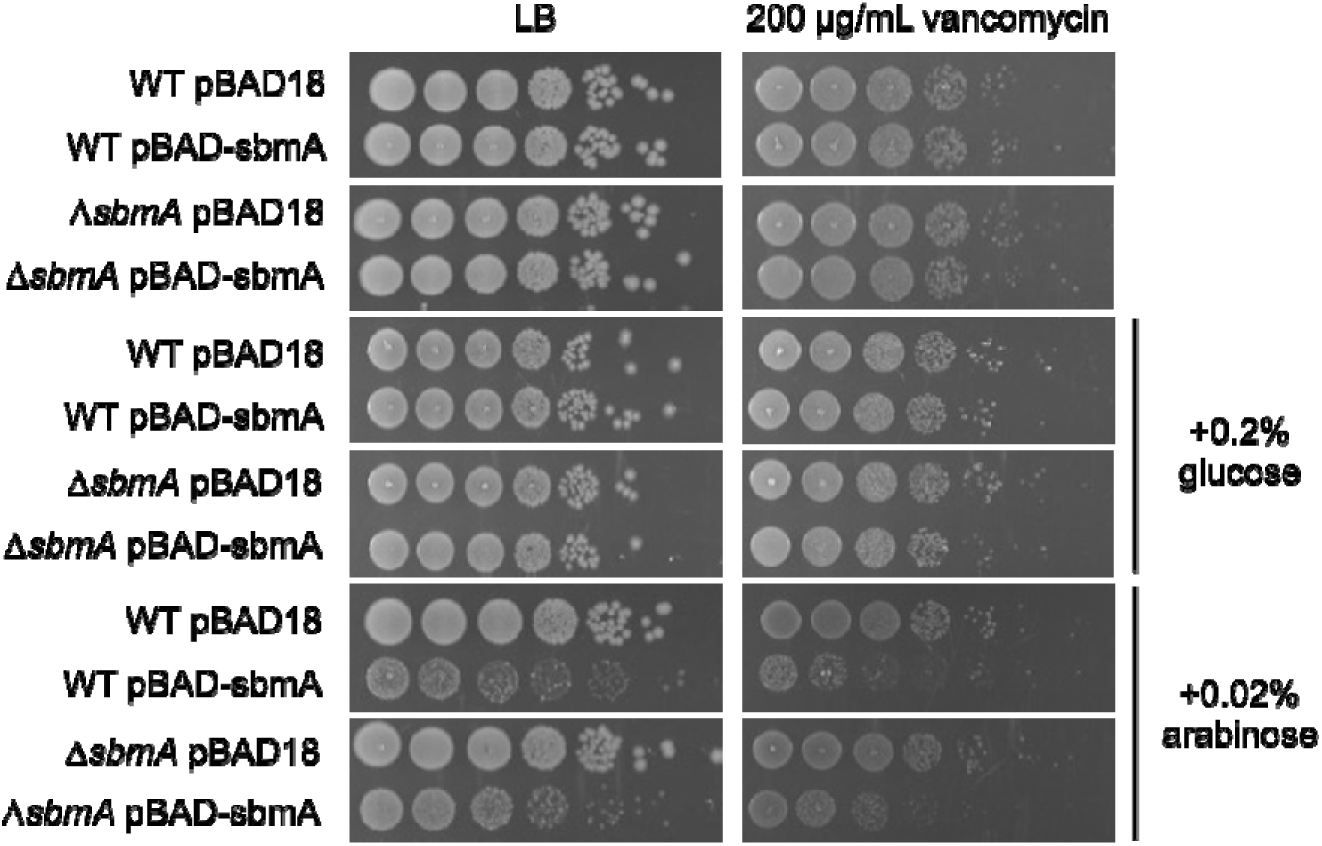
Ectopic overexpression of *sbmA* does not result in membrane envelope defects. *E. coli* BW25113 WT or Δ*sbmA* mutants transformed with a plasmid encoding for an arabinose-inducible copy of *sbmA* (pBAD-sbmA). Overnight cultures of the respective strains were diluted to OD_600_ = 0.1, ten-fold serially diluted and spotted onto LB agar + carbenicillin plates supplemented with vancomycin, arabinose and/or glucose.

